# AMLdb: A comprehensive multi-omics platform to understand the pathogenesis and discover biomarkers for acute myeloid leukemia

**DOI:** 10.1101/2023.05.19.541403

**Authors:** Keerthana Vinod Kumar, Ambuj Kumar, Kavita Kundal, Avik Sengupta, Kunjulakshmi R, Mayilaadumveettil Nishana, Rahul Kumar

**Affiliations:** Department of Biotechnology, Indian Institute of Technology Hyderabad, Kandi, Telangana 502284, India; Department of Biological Sciences, Indian Institute of Science Education and Research, Berhampur, Odisha 760010, India; School of Biology, Indian Institute of Science Education and Research, Thiruvananthapuram, Maruthamala P. O, Vithura, Kerala 695551, India

**Author notes:** Equal contribution (Keerthana Vinod Kumar, Ambuj Kumar).

**Keywords:** Acute myeloid leukemia, database, next-generation sequencing (NGS), expression, methylation, biomarkers, drug targets, mutation

## Abstract

Acute myeloid leukemia (AML) is one of the leading leukemic malignancies in adults. The heterogeneity of the disease makes the diagnosis and treatment extremely difficult. Despite the significant developments and the rapid advancements in finding new medication targets, the treatment for AML remains a challenge. With the advent of next-generation sequencing (NGS) technologies, exploration at the molecular level for the identification of biomarkers and drug targets has been the main focus for the researchers to come up with novel therapies for better prognosis and survival outcomes of AML patients. However, the massive amounts of data generated from the NGS platforms demands the necessity to create a comprehensive platform on AML to save the time invested in mining literature. To facilitate this, we developed AMLdb, an interactive multi-omics platform that allows users to query, visualize, retrieve and analyze AML-related multi-omics data. It provides a diverse collection of data resourced from various repositories allowing for a more comprehensive analysis. AMLdb contains 86 datasets for gene expression profiles, 15 datasets for methylation profiles, CRISPR-Cas9 knockout screens of 26 AML cell lines, sensitivity of 26 AML cell lines to 288 drugs, mutations in 41 unique genes in 23 AML cell lines and information on 27 experimentally validated biomarkers. The data provided can be used for deriving conclusions on potential targets for therapies, sensitivity of these targets towards the drugs, patient classification, prediction of treatment strategies and outcomes, chances of relapse etc. In this study, we have reported five genes i.e., *CBFB, ENO1, IMPDH2, SEPHS2 and MYH9* identified via our analysis using AMLdb as potential targets. Amongst this, *CBFB, IMPDH2, SEPHS2 and MYH9* have been previously validated as targets by experimental studies which is in par with our results. However, *ENO1* is a novel target identified using AMLdb which needs further investigation. We anticipates that, AMLdb can be a valuable resource to aid the research community accelerate the development of effective therapies for AML. AMLdb is freely accessible at https://project.iith.ac.in/cgntlab/amldb/ without any restrictions.

## 1. Introduction

Acute myeloid leukemia (AML) is the most common type of leukemia in adults, accounting for approximately 80% of all cases [1]. The rapid and uncontrolled growth of immature white blood cells, known as myeloblasts, in the bone marrow characterizes AML, a life-threatening malignancy that affects the bone marrow and blood [1]. The accumulated blast cells replace the normal hematopoietic cells [1]. Therefore, more than 20% of blasts in bone marrow characterize as AML [2]. It is a biologically complex disease with a high incidence and death rate worldwide [2].

According to recent studies, the total number of AML cases reported between 1990 and 2019 is 16,328,147 with total death cases accounting for 8,769,413 in the US alone [3]. Age-standardized mortality rates (ASMR) increased by 11.7% in males and 1.5% in females and age-standardized incidence rate (ASIR) increased by 34.6% in males and 7.9% in females globally [3]. In India, the incidence of AML is estimated to be around 8,000-10,000 new cases per year [4]. It is more common as people age, with cases rising from 1.3 per 100,000 in patients under 65 to 12.2 per 100,000 people in patients over 65 years of age [4].

AML is currently treated in about 35–40% of individuals under the age of 60 years, due to advancements in treatment regimens and supportive care (such as anti-infective medicines and transfusion assistance) [5]. Younger AML patients who received aggressive chemotherapy, particularly the “3 + 7 regimen” (3 days of daunorubicin + 7 days of cytarabine), which has been the mainstay of care since the 1970s, experienced long-term cures of 30 to 40% [5]. The prognosis is very dismal for those older than 60 years; most patients eventually relapse and die from their illness within a year of diagnosis [5]. Being a highly heterogeneous disease with a combination of several clinical, molecular and cytogenetic subtypes, AML demands specific targeted therapies [6]. Therefore, recent studies focus on unravelling these heterogeneities to discover directed therapies for AML subsets for better prognosis [6].

Unlike solid tumours, there are limited genes in AML genomes with a mutation frequency > 5%, e.g., *FLT3, NRAS, DNMT3A, TET2, NPM1, RUNX1, WT1, TP53, CEBPA, IDH1* and *IDH2* [7]. Utilizing next-generation sequencing (NGS), numerous genes with diagnostic, prognostic and therapeutic implications can now be evaluated simultaneously [8]. These sequencing technologies render a tremendous amount of genomic data of AML patients. This data is scattered across publicly available repositories such as The Cancer Genome Atlas (TCGA) [9], International Cancer Genome Consortium (ICGC) Data Portal [10], Gene Expression Omnibus (GEO) [11], Genomic Data Commons (GDC) data portal [12] etc. which makes it difficult for researchers to integrate and test biological hypothesis. Therefore, there is an unmet need for the systematic collection, processing, and integrated analysis of these scattered multi-omics data and the subsequent development of a user-friendly database that would shed light on the vulnerabilities of AML to design better-targeted therapies. To date, there are no comprehensive databases reported for AML that serves this purpose. Thus, we developed AMLdb, an integrative and interactive multi-omics database. AMLdb is a powerful and integrative tool with credible content that provides multi-omics information at one place. Users can explore gene expression and methylation profiles, genome-wide CRISPR-Cas9 knockout screens for various AML cell lines along with drug sensitivity. We anticipate that AMLdb will augment the ongoing effort to understanding the pathophysiology of AML. AMLdb can be accessed at URL project.iith.ac.in/cgntlab/amldb without any restriction.

## 2. Materials and Methods

### 2.1 Data Collection and Processing

The expression and methylation profiling data were searched in NCBI Gene Expression Omnibus (GEO) Database [11] by querying the keyword “acute myeloid leukemia”. We obtained 40017 results for GEO datasets and 1350899 for GEO profiles as of January 2023. Figure 1 depicts the overall workflow of AMLdb.

**Figure 1:**
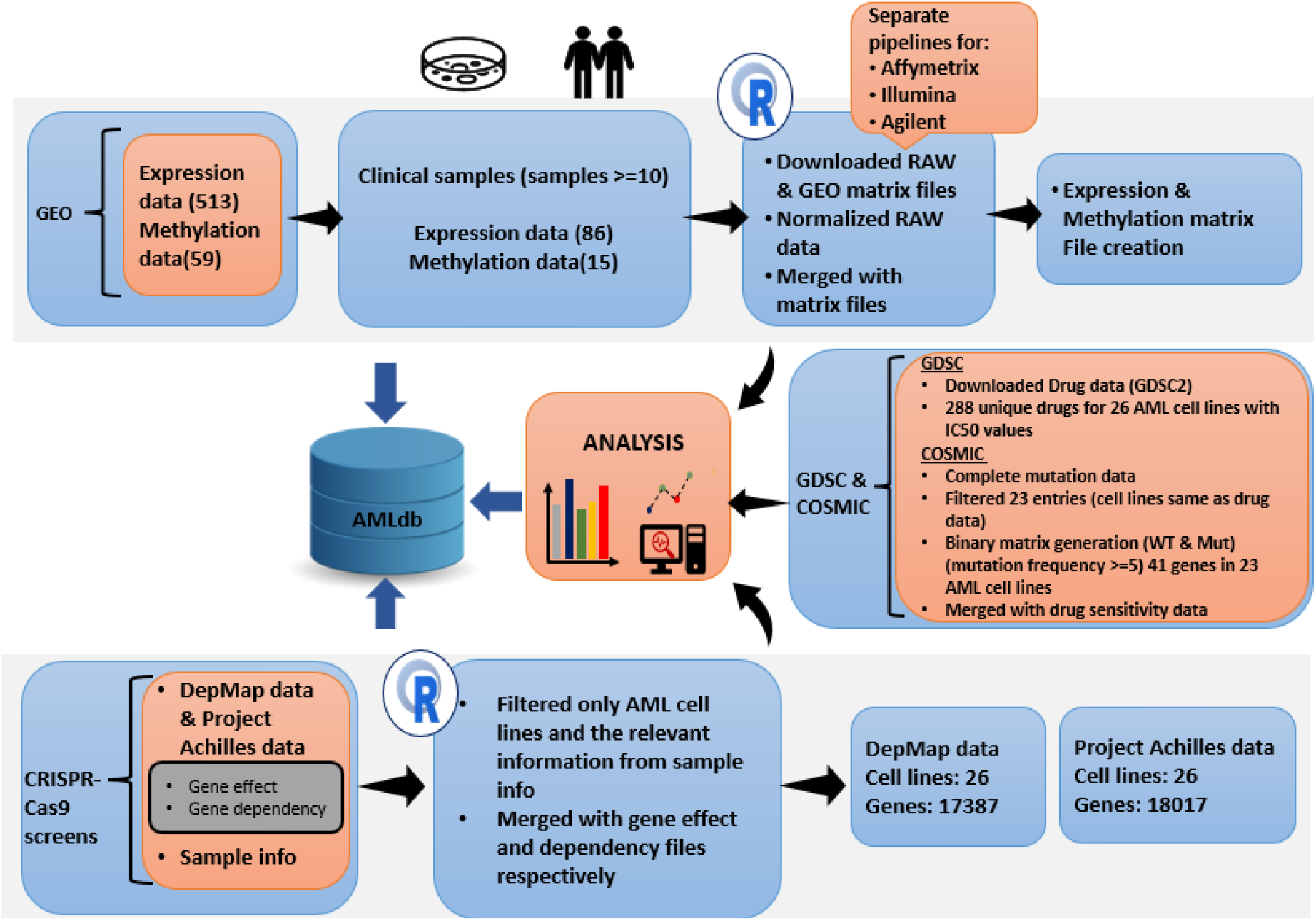
Workflow of AMLdb

#### 2.1.1 Gene expression profiling data

We searched the relevant datasets by following criteria: (a) study type: ‘Expression profiling by array’ and ‘Expression profiling by high-throughput sequencing’ (b) organism: ‘*Homo sapiens’* and obtained 513 expression profiles. We restricted the data collection process to only studies involving clinical samples with a minimum sample size of 10 and eliminated those studies that included only cell lines. With these criteria we obtained total of 485 datasets. Those datasets designated as sub-series were included while excluding their super-series to prevent duplication. Any datasets with truncated or corrupted data (e.g., some datasets did not contain both Gene symbols and Entrez IDs) were not included. We did the manual review and curation of remaining datasets by carefully removing the normal samples and retaining only the tumour ones ensuring they have all relevant information. After applying these criteria, we finally obtained 86 datasets for expression profiling with 5674 tumour samples (including both “by array” and “by high-throughput sequencing”).

#### 2.1.2 Gene methylation profiling data

We used the following criteria for searching the methylation profiles in GEO: (a) study type: ‘Methylation profiling by array,’ ‘Methylation profiling by SNP array,’ and ‘Methylation profiling by high-throughput sequencing’ (b) organism: *Homo sapiens* and obtained 135 methylation profiles. We further select the studies with sample size >= 10 and obtained 59 methylation profiles. We reviewed these datasets individually to eliminate all the truncated and/or corrupted data and finally obtained 15 datasets for methylation profiling with 874 tumour samples.

#### 2.1.3 CRISPR-Ca9 knockout screens

To identify crucial genes responsible for the survival and proliferation of cancer cells in AML, we downloaded data from CRISPR-Cas9 based knockout screens from the DepMap portal (version: 22Q2) [13]. We filtered the sample information data to obtain all relevant information like DepMap ID, cell line names, sample collection site, primary/metastasis, lineage, sex, and age, and merged it with the gene effect scores. Then, we created a final table containing gene effect scores of 17387 genes in 26 different AML cell lines and stored it in AMLdb.

We also downloaded and used gene effect scores from Achilles project available at the DepMap portal (version: 22Q2) [13], which utilizes CRISPR-Cas9 mediated knockout screens to identify genes essential for the survival of cancer cells. We used a similar pipeline to process data and generate the final data table for Achilles project containing gene effect scores of 18017 genes in 26 different AML cell lines.

#### 2.1.4 Drug sensitivity data

We downloaded the drug sensitivity data from the Genomics of Drug Sensitivity in Cancer (GDSC) portal [14] to understand the sensitivity of AML cell lines to anticancer drugs. To do this, we selected “GDSC2” screening set, “all” as the target pathway, and “Acute Myeloid Leukemia” as the tissue under the drug data category. The downloaded file contained 288 unique drugs for 26 unique AML cell lines and their IC50 values.

#### 2.1.5 Mutation data

To incorporate the mutation data, we obtained the complete mutation data from the Catalogue Of Somatic Mutations In Cancer (COSMIC) database [15] by accessing downloads section (version v97 released on 29-NOV-22). We got a table of 6057239 entries. From this, we filtered “Acute Myeloid Leukemia” from “Histology subtype 1” category, which resulted in 119636 entries and selected only those entries having the same cell lines as drug sensitivity data. Finally, we removed those entries wherever “Mutation description” was “Unknown”, “Substitution - coding silent,” and “Nonstop extension” rendering a table of 7250 unique genes along with their mutation details in 23 AML cell lines.

### 2.2 Database Design and Implementation

AMLdb is a user-friendly web database that utilizes MySQL/10.4.25-MariaDB as its Database Management System (DBMS). We used HTML5, JavaScript, and PHP5 to develop the web interface to provide structure and dynamicity, while CSS3 enhanced its visual appearance and design. HTML Codex provided the template design [16] and we constructed the database on an Ubuntu-based system utilizing the Apache HTTP Server (v2.4.54). Furthermore, it is compatible with commonly used browsers such as Chrome, Firefox, Internet Explorer and Safari.

## 3. Results

### 3.1. Database Statistics

AMLdb is a comprehensive platform containing 101 datasets; 86 for expression profiles with 5674 samples and 15 for methylation profiles with 874 samples. We have resourced 26 AML cell lines with their corresponding CRISPR-Cas9 knockout screens data from DepMap (17387 genes) and Achilles project (18017 genes). Information on 288 anticancer drugs and their effect on 26 AML cell lines has been resourced from the GDSC database. AMLdb also contains mutation data of 23 AML cell lines from the COSMIC database. Figure 2 describes the user-interface and statistics of AMLdb.

**Figure 2:**
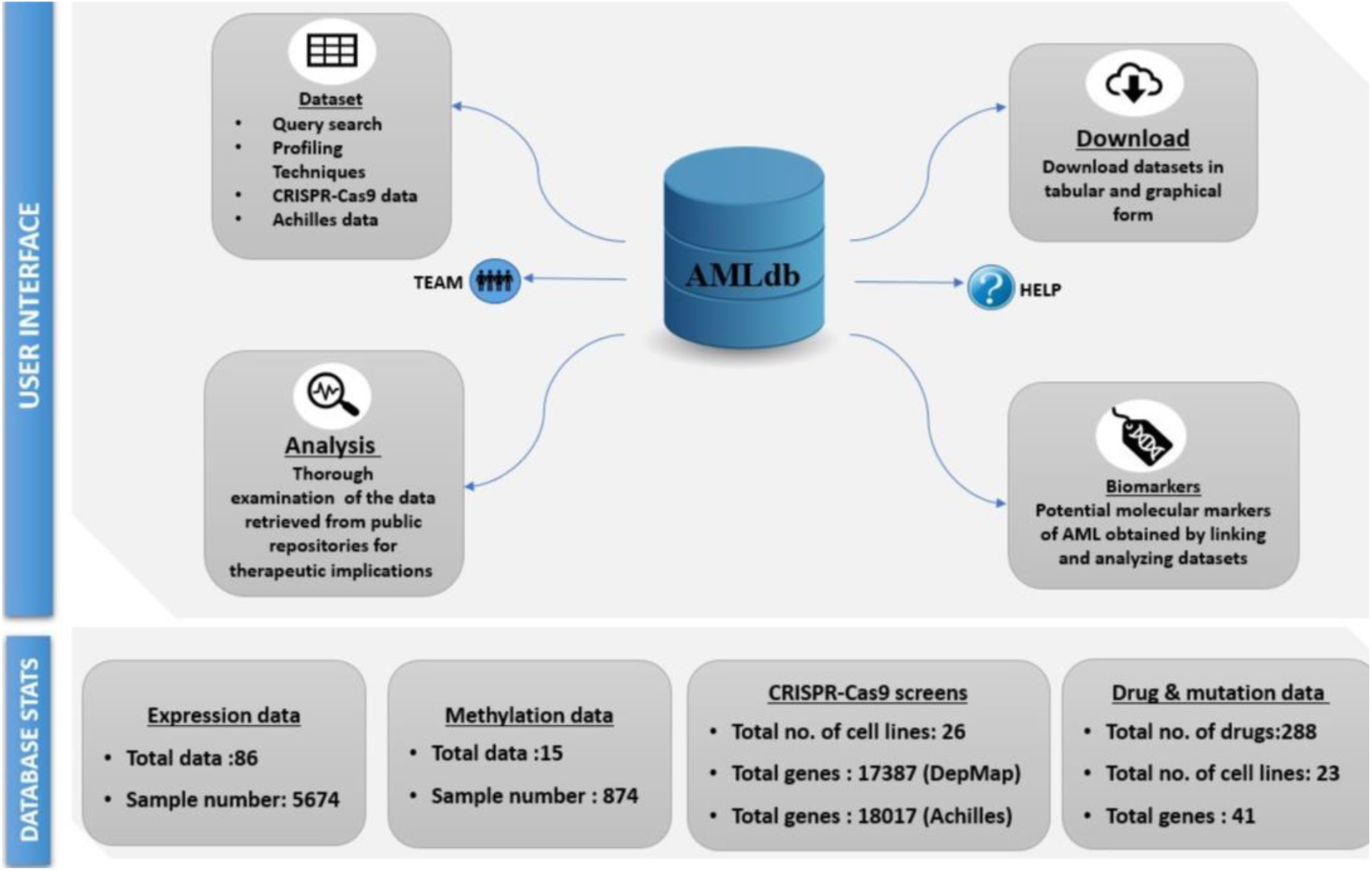
User interface and statistics of AMLdb

### 3.2 Database Content and Web Interface

We have developed AMLdb as a multi-omics platform with a user-friendly interface for querying, visualizing and retrieving data related to AML. It offers seven sections to the users in its navigation bar: Home, Dataset, Analysis, Biomarker, Download, Help and Creators.

#### 3.2.1 Home page

It is the introductory start-up page for the website, which displays information about the database contents and shows all the options available for search for the user. Figure 3 shows the Home page of AMLdb.

**Figure 3:**
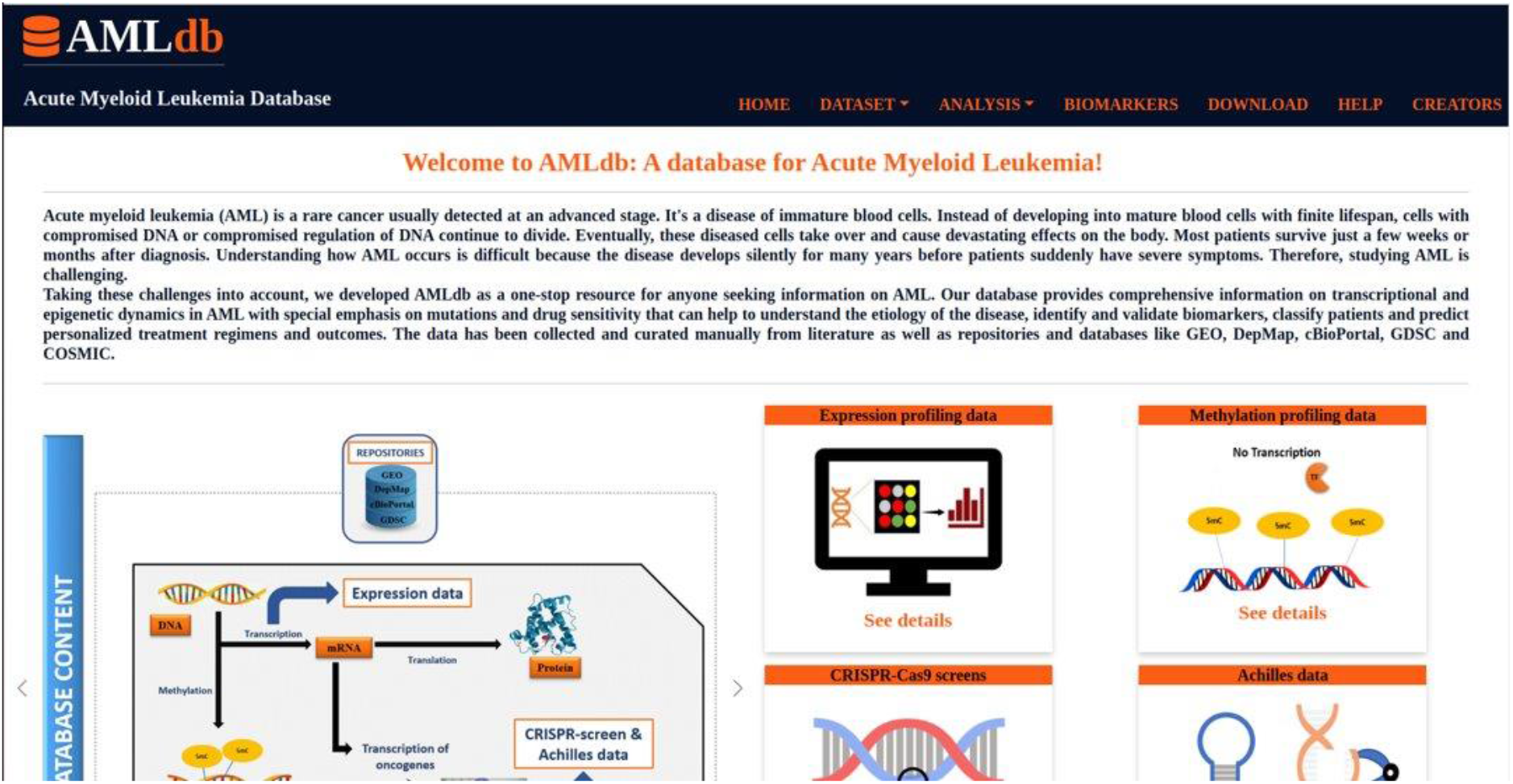
Home page of AMLdb along with the navigation bar showing the options Home, Dataset, Analysis, Biomarker, Download, Help and Creators.

#### 3.2.2 Dataset page

This has three modules: ‘GEO,’ ‘CRISPR-Cas9 screens’ and ‘GDSC.’ Under ‘GEO,’ users can enter their query from the Query search category (expression or methylation data) or Profiling Technique. Query search category provides a form consisting of six options: Dataset ID, PubMed ID, profiling technique, processing package, sample source and type of data. Upon submission, a table of queried search containing information like Dataset ID, PubMed ID, Platform, sample number, sample source, profiling technique, genes, probes, etc., would appear. A download option is made available for downloading the data in the csv format. We linked the GEO ID’s and PubMed ID’s to their respective sources. The second option under ‘GEO,’ ‘Profiling Techniques’ will allow users to browse the datasets based on sequencing platforms such as Affymetrix, Illumina and Agilent.

The ‘CRISPR-Cas9 screens’ module consists of two options; ‘DepMap’ and ‘Project Achilles’. With these options, users can search for any gene for their dependency scores in 26 AML cell lines along with clinical information. The “Click to view plot” button provides a plot of cell lines on X axis with Gene effect on Y-axis. The ‘GDSC’ module contains the “Drug Search” option where the user can select a drug name from the dropdown menu which provides a table of cell lines and the respective logIC50 values. The “Click to view the plot” option provides a graph of cell lines with logIC50 on Y-axis of selected drug in 23 AML cell lines (excluding those with null values). These plots have been made using CanvasJS [17].

#### 3.2.3 Analysis page

This page has two modules: ‘Profiling’ and ‘Drug sensitivity’. The ‘Profiling’ module contains two options: Expression and Methylation. In both options, the user can query any gene in AMLdb, gene expression of queried gene will be displayed as Box-Whisker’s plots. Each boxplot shows the mean expression values of the queried gene in Affymetrix, Illumina, and Agilent platforms. Five boxplots are shown in the database, as the expression values for the genes have been processed by using various processing methods like Robust Multi-Array (RMA) and background correction and quantile normalization for Affymetrix, quantile normalization and transcripts per million (TPM) for Illumina and background correction and quantile normalization for Agilent. The ‘Drug Sensitivity’ module includes mutational analysis. Here, the user can select a particular gene and a drug from the dropdown menu to retrieve a table of gene name, drug name, names of mutated and wild-type cell lines along with their corresponding logIC50 values and the p-value. With this option, user can check the effect of gene mutation on drug sensitivity. Further, “Click to view plot” button provides a Box-Whiskers plot showing mutational analysis of the Gene (wild type & mutated) vs logIC50 values and their p-value.

#### 3.2.4 Biomarker section

This page provides tabulated information of 27 potential AML biomarkers with information like biomolecule, regulation status, type, subject, experiment, significant values etc.

#### 3.2.5 Download section

This page allows users to directly download individual datasets uploaded on the website.

#### 3.2.6 Help section

This page will provide a detailed manual of the website for the user’s easy access.

#### 3.2.7 Creators section

This page provides a comprehensive overview and details of the individuals involved in creating and maintaining the database, including their names, roles, and contact information.

## 4. Integrated analysis with AMLdb

### 4.1 Impact of gene mutation on drug sensitivity

From the 7250 unique genes obtained from COSMIC database, we generated a matrix of all the genes with their mutation status in 23 AML cell lines. We selected only those genes with a mutation frequency greater than or equal to 5 (n=41). We conducted the mutational analysis of these 41 genes, across 23 AML cell lines by integrating this information with the corresponding drug sensitivity profiles of 288 drugs from GDSC. Subsequently, we conducted t-tests for statistical analysis. We also generated Box-Whisker plots for each queried gene, displaying the logIC50 values for mutated and wild type AML cell lines for that gene. To show the utility of this tool, we selected two associations i.e., *AHNAK2*-Nutlin-3a (-) and *NRAS*-Entospletinib, which shows gene mutations decreasing the sensitivity of AML cell lines towards respective anticancer drugs (Figure 4A and 4B). In two other associations, *BIRC6-*Foretinib and *KMT2C-*Methotrexate, gene mutations sensitising the AML cell lines towards respective anticancer drugs has been shown (Figure 4C and 4D).

**Figure 4:**
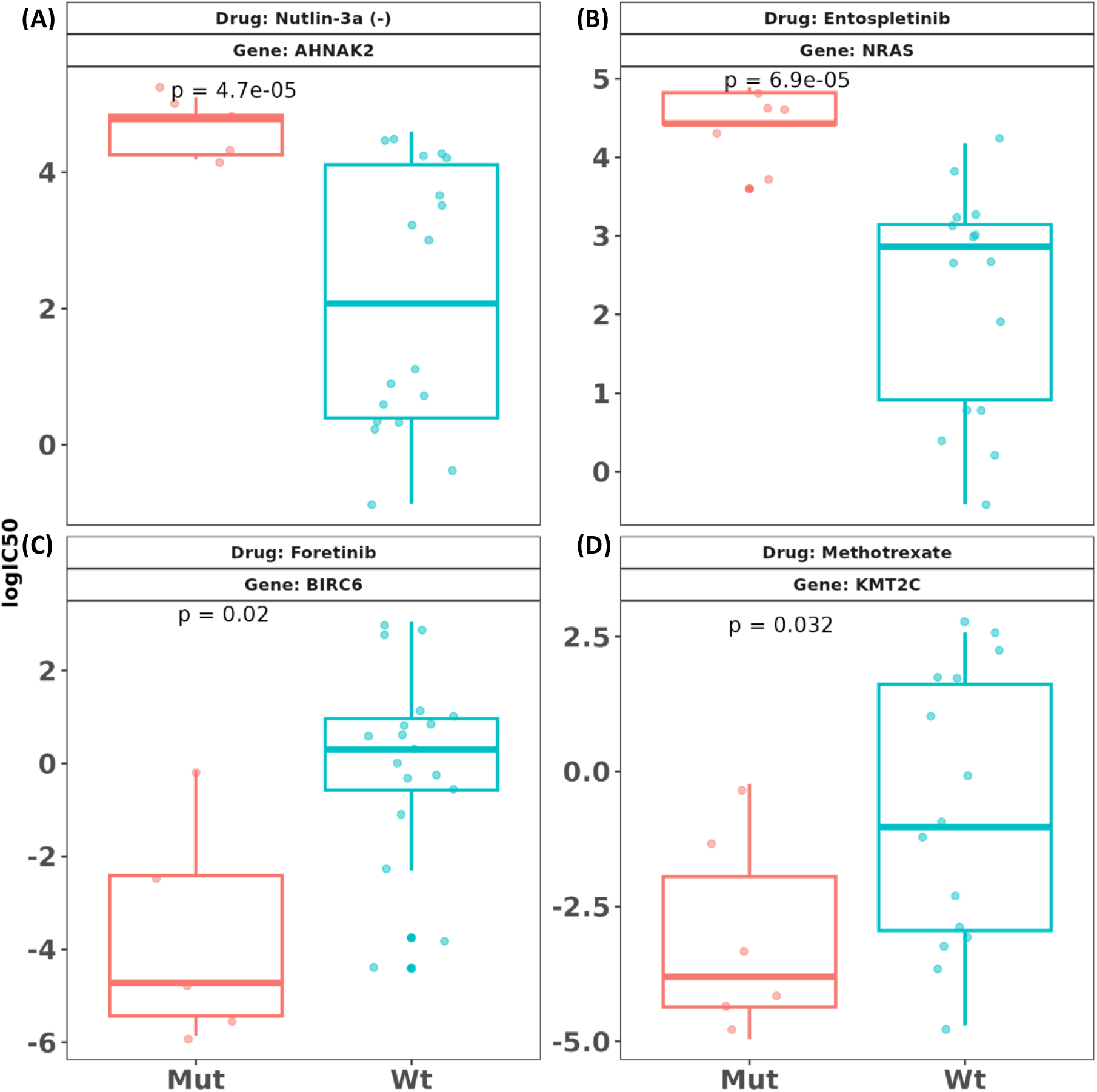
Mutation in genes (A) *AHNAK2* and (B) *NRAS* decrease the sensitivity of drugs Nutlin-3a (-) and Entospletinib for AML cell lines respectively. Mutation in genes (C) *BIRC6* and (D) *KMT2C* sensitize AML cell lines for the drugs Foretinib and Methotrexate respectively.

### 4.2 Novel gene targets for AML

We performed an integrated analysis using all the Robust Multi-Array (RMA) and quantile normalized expression datasets of Affymetrix platform. For each dataset, we generated the top quantile genes (assuming them as over-expressed in AML) after taking the mean expression values of each gene across all the tumour samples. We removed all the house-keeping genes [18] and merged them with the gene dependency scores from CRISPR-Cas9 knockout screens. We then counted for the number of cell lines where dependency scores were more than 0.7. We filtered the genes with counts >=13 (i.e., considering that fact that 50% of the AML cell lines were dependent on those genes for their survival). We performed this pipeline for all the datasets and filtered the common genes. Thus, we got 37 genes common for Affymetrix RMA data and 45 genes for quantile normalized data. Further, we eliminated the genes which are common across different cancer types as provided in DepMap portal. As a result, we got five genes exclusive for AML; *CBFB, ENO1, IMPDH2, SEPHS2* and *MYH9*. Literature survey on these genes suggest the direct link of *CBFB, IMPDH2, SEPHS2* and *MYH9* with AML pathology [19–22]. Our analysis suggests further investigation for delineating the link of *ENO1* with AML pathology.

## 5. Conclusion

The AMLdb is currently the only database that consolidates information about AML-specific gene expression, methylation, drug sensitivity, gene essentiality and biomarkers data from published literature, along with their experimental results, in a single location. Figure 5 depicts the significance of AMLdb. AMLdb eases the tedious task of streamlining the vast amounts of multi-omics data that are widely scattered across different platforms to produce conclusions on potential biomarkers and drug targets. All the data uploaded has been uniformly processed and curated manually to maintain the homogeneity. AMLdb provides an interactive user-interface and allows users with minimum or no knowledge of computer programming or bioinformatics skills to analyse and retrieve the data without any restriction. We anticipate that AMLdb will be a one-point solution for the research community to analyse multi-omics data on AML.

**Figure 5:**
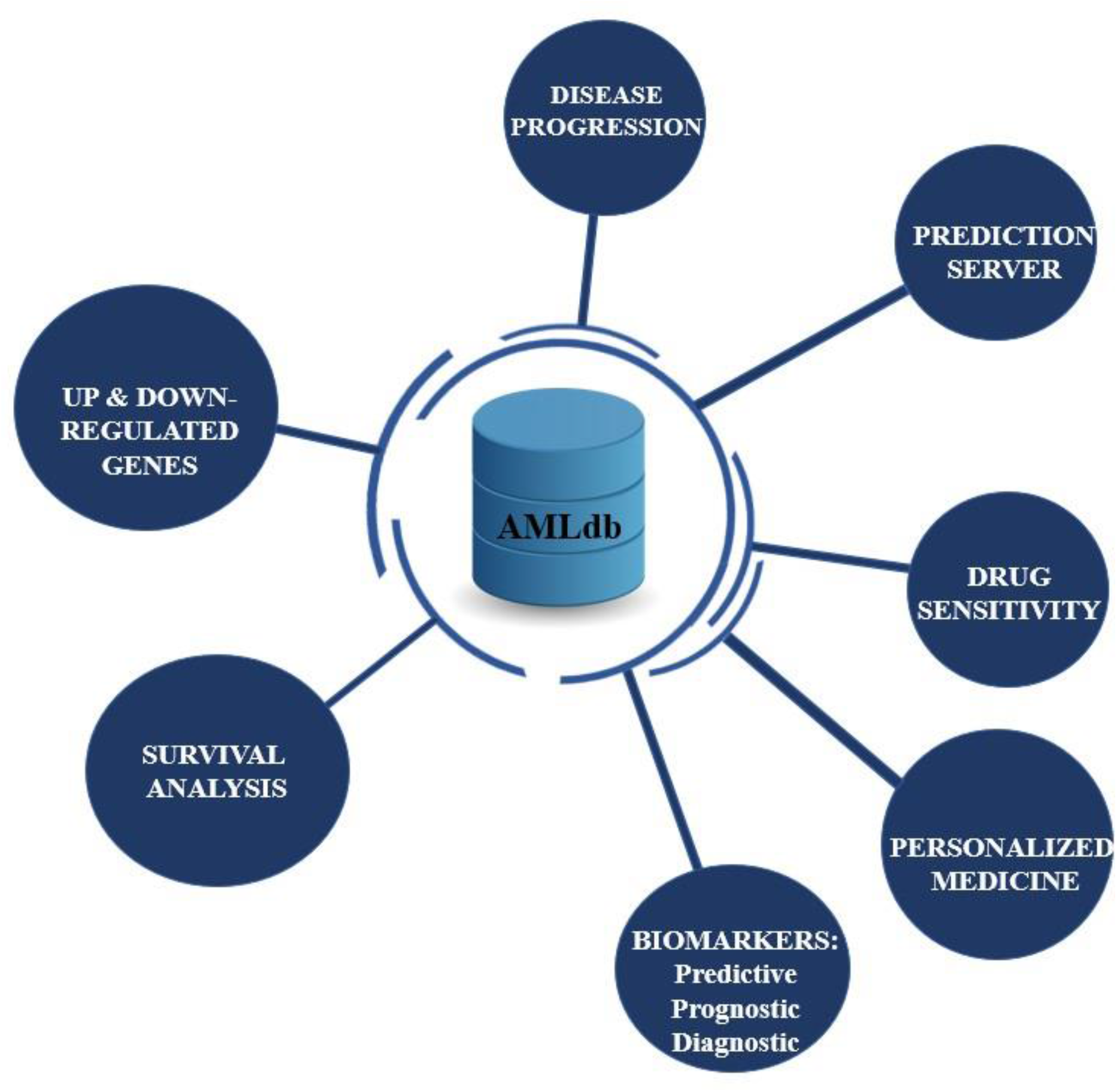
Significance of AMLdb and the potential inferences that can be obtained from it

## 6. Discussion

AMLdb will play a crucial role in advancing our understanding of AML pathology and it will have vast implications for disease progression, survival analysis, and personalized medicine. One key area where AMLdb excels is in the identification of up-regulated and down-regulated genes associated with AML. By analysing gene expression and methylation profiles, AMLdb facilitates the identification of genes that are dysregulated in AML patients. This information is vital for understanding the underlying molecular mechanisms of the disease and can guide further research into targeted therapies. Disease progression is a crucial aspect of AML research. By integrating clinical data with molecular and genetic information, AMLdb enables researchers and clinicians to identify prognostic factors that influence patient outcomes. This knowledge can aid in making informed decisions regarding treatment strategies and patient management. AMLdb incorporates information on drug sensitivity profiles that can aid in the selection of appropriate personalized therapies, improving treatment efficacy and reducing the risk of drug resistance. It also serves as a rich source for biomarker discovery. The database facilitates the identification of predictive, prognostic, and diagnostic biomarkers that can assist in early detection, risk stratification, and monitoring of AML patients. These biomarkers hold significant potential for guiding clinical decision-making, improving patient outcomes, and fostering the development of novel therapeutic approaches. AMLdb’s design allows for the possibility of developing a prediction server utilizing the data uploaded into the database. The incorporation of a prediction server into AMLdb would enable researchers to identify specific molecular markers or genetic alterations that are predictive of treatment response or disease progression.

## 7. Author Contribution

KVK, AK and RK conceived the idea and prepared the database framework. KVK and AK collected and analysed the datasets. KVK developed the database and wrote the first draft of the manuscript. KK, AS and KR helped in collecting the datasets and preparing the database outline. All authors read the manuscript and approved it for submission.

## 9. Acknowledgements

KVK, AK, and KK acknowledge the fellowship from Ministry of Education, Government of India. AS acknowledge the fellowship from Council of Scientific and Industrial Research (CSIR), Government of India. We are thankful to the Indian Institute of Technology Hyderabad for providing research infrastructure.

